# Estimating the Biological Validity of the DSM for Attention Deficit/Hyperactivity Disorder Using Multivariate Analysis for Small Samples

**DOI:** 10.1101/126433

**Authors:** Dimitri M. Abramov, Evelyne Vigneau, Saint-Clair Gomes-Junior, Carlos Alberto Mourão-Júnior, Monique Castro-Pontes, Carla Quero Cunha, Leonardo C. deAzevedo, Vladimir V. Lazarev

## Abstract

**Background.:** Psychiatric nosology lacks objective biological foundation, as well as typical biomarkers for diagnoses, which raises questions about its validity. The problem is particularly evident concerning Attention Deficit/Hyperactivity Disorder (ADHD). The objective of this study is to estimate whether the “Diagnostic and Statistical Manual of Mental Disorders” (DSM) is biologically valid for ADHD diagnosis using a multivariate analysis for small samples from a large dataset concerning neurophysiological, behavioral, and psychological variables.

**Methods:** Twenty typically developing boys and 19 boys diagnosed with ADHD, aged 10-13 years, were examined using the Attentional Network Test (ANT) with records of event-related potentials (ERPs). From 815 variables, a reduced number of latent variables (LVs) were extracted with a clustering method, for further reclassification of subjects using the k-means method. This approach allowed multivariate analysis to be applied to a significantly larger number of variables than the number of cases (E. Wigneau et al., 2003, 2015)

**Results:** From datasets including ERPs from the mid-frontal, mid-parietal, right frontal, and central channels, only seven subjects were miss-reclassified by the LVs. An estimated specificity of 75.00% and sensitivity of 89.47% for DSM were found in the reclassification. The kappa index between DSM and behavioral/psychological/neurophysiological data was 0.75, which is regarded as a “substantial level of agreement”.

**Discussion:** Results showed that CLV is a useful method for diagnostic classification using a large dataset of small samples, suggesting the biological validity of DSM for ADHD diagnosis, in accordance to alterations in fronto-striatal networks previously related to ADHD.

## Introduction

The validity of the *Diagnostic and Statistical Manual of Mental Disorders* ‐DSM (American Psychiatric Association, 1994 – 2013) [1,2] is usually questioned due to its subjective criteria and to the absence of objective tests to define nosological entities [3, 4]. Therefore, the lack of objective determinants makes room for criticism and raises doubts about the nosological criteria itself [4]. This is particularly important in controversial mental diseases such as the Attentional Deficit/Hyperactivity Disorder – ADHD [5,6]. Similar to some other disorders, ADHD diagnosis is based on the quantification of normal behavior characteristics using DSM scores. Thus, DSM has a taxonomy of categories that are defined by dimensional phenomena based on a variation from normality [7,8]. Correlations between biological alterations and ADHD are still not enough to support the diagnosis [8, 9].

Mental disorders are multidimensional complex entities that can be mostly associated with a pool of biological and neurobehavioral variables, often with non-linear relationships [8, 10], rather than being defined by a single distinct biomarker such as an antibody or aberrant protein [11]. This formally determines their complexity; a complex nosological entity such as ADHD may be defined by a large set of different quantitative variables. From this standpoint, multivariate factor analysis techniques have been applied to electroencephalographic or biochemical data since the 1970’s in order to develop high-sensitivity and accuracy models for the diagnosis of mental disorders [12-16]. However, in order to explain these complex mental conditions in terms of biological correlates, a substantially larger sample of subjects than those set of variables is required. Thus, high-dimensional datasets from relatively few observations or subjects usually appear to be statistically questionable and hamper unambiguous interpretation. In order to tackle this problem (the curse of the dimensionality), approaches based on feature selection in a supervised context or on feature extraction in explorative data analysis (for instance, Principal Components Analysis, Canonical Correlation Analysis) have been investigated. Particularly, a strategy of unsupervised “clustering of variables around latent variables” (CLV) was used, which makes it possible to extract synthetic/latent variables through cluster analyses [17-19]. These few latent variables represent dataset informativeness for a classifier methodology in a subsequent step (e.g. inputs for a k-means clustering algorithm). This was the first attempt to apply the CLV method to a large psychophysiological dataset. This method is expected to help solve an important methodological problem of multidimensional approaches in clinical research when recruiting large groups of patients is difficult.

The present study is an attempt to discuss whether DSM is valid for an accurate diagnosis of the nosological entity called ADHD, analyzing a small sample of subjects with well-controlled confounding variables and bias. Large datasets of (1) behavioral variables, (2) neurophysiological data (event-related potentials – ERPs, including late cognitive component, P3 or P300 [20,21]), both obtained while subjects performed the Attention Network Test - ANT [22,23], as well as (3) psychological characteristics (from WISC-III) have been collected from typically developing boys (TD) and boys diagnosed with ADHD using the DSM-IV-TR criteria.

The relationships between ADHD manifestations, evaluated using DSM-IV-TR, and the objective characteristics obtained by the above-mentioned experimental measurements were here analyzed. The aim was to discuss the taxonomic (i.e. classificatory) aspects of ADHD diagnosis using DSM, compared with a data-driven statistical approach, without considering the role of these characteristics in putative brain mechanisms underlying ADHD. This study does not address controversial issues, e.g. if the ADHD phenomenology actually constitutes a mental disorder or not. The present study seeks to discuss the validity of the DSM as a tool to identify this phenomenology, making inferences about its sensitivity and specificity.

## Methods

### Design and Subject Selection

This transversal and exploratory study was conducted in accordance with the Declaration of Helsinki and approved by the Ethics Committee of the National Institute of Women, Children, and Adolescents Health Fernandes Figueira (CAAE 08340212.5.0000.5269). All the participants provided their oral consent in the presence of their caregivers, who provided written informed consent after receiving a complete description of the study.

Sixty boys, aged 10-13 years, were included according to DSM-IV-TR: 35 with ADHD (coded as T001 to T035) and 25 TD (coded as C001 to C025). The exclusion criteria were: (1) history of chronic diseases, and any suspicion of psychiatric disorders other than ADHD (psychosis, affective, obsessive-compulsive and tic disorders, phobic and post-traumatic stress conditions, anorexia, bulimia, encopresis, or enuresis) as screened by K-SADS-PL [23]; (2) use of any psychotropic medicines for at least 30 days; (3) estimated I.Q. equal or lower than 80; and (4) less than 6 hours of regular sleep and (5) report of somnolence before the ANT testing.

The following were considered as confounding variables: years in school, monthly family income (in Brazilian Reals), sleep hours the night before, and mean weekly time which the boy dedicated to computer activities and videogames (ranked as follows: 1, less than 2h/week; 2, 2-4h; 3, 5-8h; 5, more than 15 h/week), learning rate (see below) in the experimental procedure, as well as age.

After applying the exclusion criteria, the behavioral analysis of accuracy (AC) and speed-accuracy tradeoff (see below), and after excluding confounding outliers, 19 ADHD and 20 TD boys remained in the sample, and their data were considered as experimental results.

### Clinical and Psychological Examination

Each subject was evaluated using a structured interview where his or her caregivers were shown the DSM-IV-TR criteria and were instructed to point out carefully whether or not each specific criterion was an exact characteristic of their child’s behavior. If there was any doubt about or hesitation in any item, it was disregarded. Thus, subjects were classified in accordance with the DSM-IV-TR.

Intelligence Quotient (I.Q.) was estimated by Block Design and Vocabulary subtests of the Wechsler Intelligence Scale for Children, WISC-III [24, 25]. The Arithmetic and Digit Spam subtests were also performed for the pool of variables used in the multivariate analysis.

### Experimental Procedures

The ANT version adapted for children was used [20, 26]. A forced, two-choice test was performed, where the subject was instructed to observe the horizontal orientation of a target stimulus (yellow fish), which was (or not) preceded by a cue signal (a red star). The target appeared above or below the fixation point. There were three equiprobable cue conditions corresponding to this signal’s position or to its non-appearance: 1) at the subsequent upper or lower position of the target - spatial cue condition; 2) at the central fixation point - neutral cue condition; or 3) no cue condition. The subject had to promptly press the left or right arrow key on the keyboard, according to the horizontal orientation of the target. The test was organized in 8 test blocks, with 24 trials each, and one preceding training block. Reaction time (RT) and accuracy (AC, rate of hits) were recorded.

Subjects with AC lower than 70% and speed-accuracy tradeoff (estimated as AC x RT) lower than the mean sample value minus two standard deviations were excluded.

### EEG Acquisition

During the ANT, the subject’s EEG was recorded using a Nihon Kohden NK1200 EEG System at 20 scalp points according to the International 10/20 System (Figure 1), with reference at the lateral central leads (linked C3 and C4, the physical reference of the System). Impedance was below 10 kΩ, sampling rate was 1000 Hz with a 16-bit resolution, and the filters were as follows: low-pass 0.5 Hz, high-pass 100 Hz, and notch 60 Hz.

**Figure 1.**
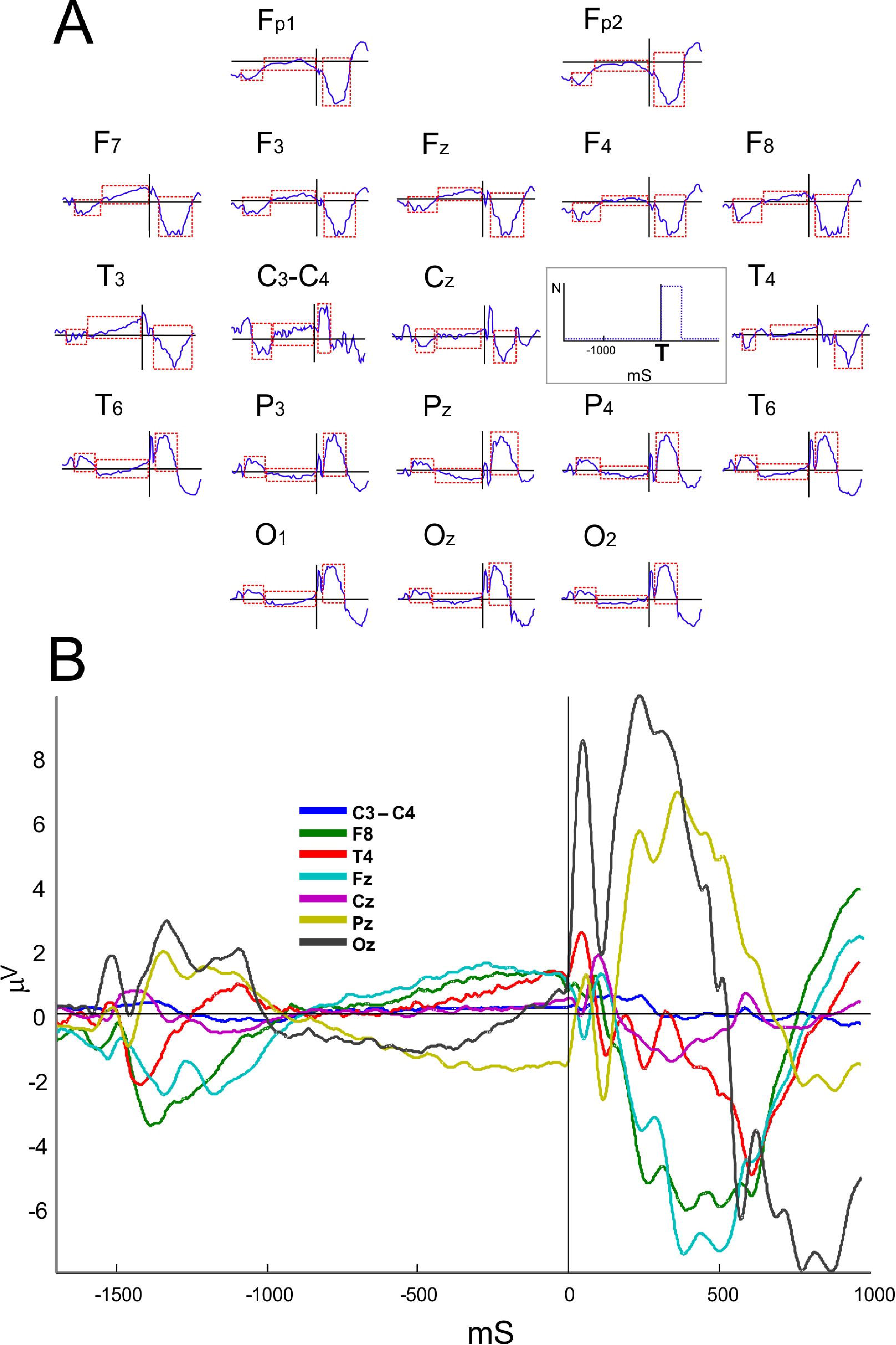
Electrical potentials related to Attention Network Test. (A) Cue‐ and target-related potentials and variation in interstimulus voltage during Attention Network Test performance (averaged over all ANT conditions and all subjects). Overview of the ERPs for all channels, including ‘C3 minus C4’ (normalized amplitudes, μV), with waves of interest (in dotted squares) to calculate the mean square of the total area under each wave and peak amplitudes/latencies. In the grey box, C and T stand for cue and target onsets, and the trigger signal is identified by dotted lines. Above the ERP graphs, scalp areas are marked with letters with even (right), odd (left) or z (midline) indices: occipital – O, parietal – P, central – C, frontal – F, frontopolar – Fp, mid-temporal – T, posterior temporal – T5 and T6, and anterior temporal – F7 and F8. Scalp scheme for 10-20 montage at the bottom left. (B) ERPs for selected channels, in different colors, showing wave amplitudes (μV).

## Data Analysis

Predictor (i.e. DSM score for ADHD as the sum for inattention and hyperactivity/impulsivity scores) and confounding variables were compared between the groups using Student’s t-test, after normality and heteroscedasticity assessments were performed using the Shapiro-Wilk and Levene’s tests, respectively (Table 1).

**Table 1.** Predictor and Confounding Variables between Gr176 oups

|  | Typically<br>Developing<br>(n = 20) |  | ADHD<br>(n = 19) |  | T-value<sup>a</sup> | p-value |
| --- | --- | --- | --- | --- | --- | --- |
|  | M | SD | M | SD |  |  |
| Predictor Variables (DSM scores) |  |  |  |  |  |  |
| **Inattention** | **2.40** | **1.66** | **7.21** | **1.27** | **-10.09** | **< 0.001** |
| **Hyperactive+Impulsive** | **2.60** | **1.63** | **4.36** | **2.90** | **-2.36** | **0.023** |
| **Total** | **5.00** | **2.75** | **11.63** | **3.00** | **-7.19** | **< 0.001** |
| Confounding Variables |  |  |  |  |  |  |
| Age (years) | 11.30 | 0.86 | 11.52 | 1.07 | -0.73 | .471 |
| **Estimated I.Q.** | **109.35** | **13.53** | **97.36** | **12.53** | **2.86** | **0.006** |
| Hours of Sleep (previous night) | 7.25 | 2.12 | 7.68 | 1.73 | -0.70 | 0.490 |
| Videogame <sup>b</sup> | 3.40 | 1.23 | 2.89 | 1.66 | 1.08 | 0.286 |
| Computer handling <sup>c</sup> | 3.30 | 1.26 | 2.94 | 1.68 | 0.74 | 0.462 |
| Years in school | 6.05 | 1.14 | 6.10 | 1.32 | -0.14 | 0.889 |
| Family Incomes <sup>d</sup> | 6355.00 | 5643.57 | 3978.94 | 4436.91 | 1.46 | 0.153 |
| Learning Index <sup>e</sup> | 1.22 | 0.34 | 1.26 | 0.30 | -0.36 | 0.721 |

<sup>a</sup> Inference using Student's t-test for independent samples, significant differences (*p* value < 0.05) in bold. <sup>b, c</sup> Videogame and computer handling were scored as: 1 = less than 2h/week; 2 = 2-4h/wk, 3 = 5-8 h/wk, 4 = 8-15 h/wk, 5 = more than 15h/wk. <sup>d</sup> in Brazilian Reals (R\$), per month <sup>e</sup> Learning Index in the ANT: reaction time (RT) (ms) of last trial block / mean RT (ms) of the first experimental blocks.

Several behavioral variables were gathered from the ANT for each subject: mean AC of task performance, i.e. the percentage of correct responses; mean Reaction Time (RT) and its standard deviation, called intra-individual variation of RT (IVRT), for each cue and target condition. Additionally, data included the ANT scores of the subjects: alerting, orienting, and conflict resolution, mean RT for correct and incorrect responses, and learning rates (the mean scores of the first ANT trial block divided by those of the last experimental block evaluated for RTs and IVRTs).

The following ERP characteristics were estimated inside time intervals of interest (see dotted squares in Figure 1 and their measures in Table S1) for each subject, in each cue and target condition and in each EEG derivation (except for C3 and C4): (1) peak amplitude, (2) peak latency, and (3) the total amplitude under the wave (i.e. the sum of all amplitude values, proportional to the mean amplitude). These three time intervals included late ERPs evoked by 1) the cue, 2) the target stimuli, and 3) the voltage variation between these two ERPs. The same ERP parameters were calculated for the difference between signals of the two central channels (C3 minus C4) instead of estimating the absolute signal values in each of these leads, observing the EEG reference used here (C3+C4).

Thus, the total number of variables of interest (VOI) was 774: four scores obtained from the above-mentioned WISC-III subtests, 29 behavioral variables obtained from ANT, and 741 neurophysiological variables (39 for each of the 18 channels and C3-C4).

The neurophysiological variables obtained represent topographical parameters of regional (local) cerebral activity, while the variables from ANT and WISC provide the integral characteristics of the subject. For the analysis, the activity in certain brain regions of the subjects performing ANT (but not all) was assumed to best correlate with ADHD phenomenology. For this reason, 40 different sets of channels (Table 2) were used for data mining, in an attempt to find the best agreement with DSM (if any).

**Table 2.**
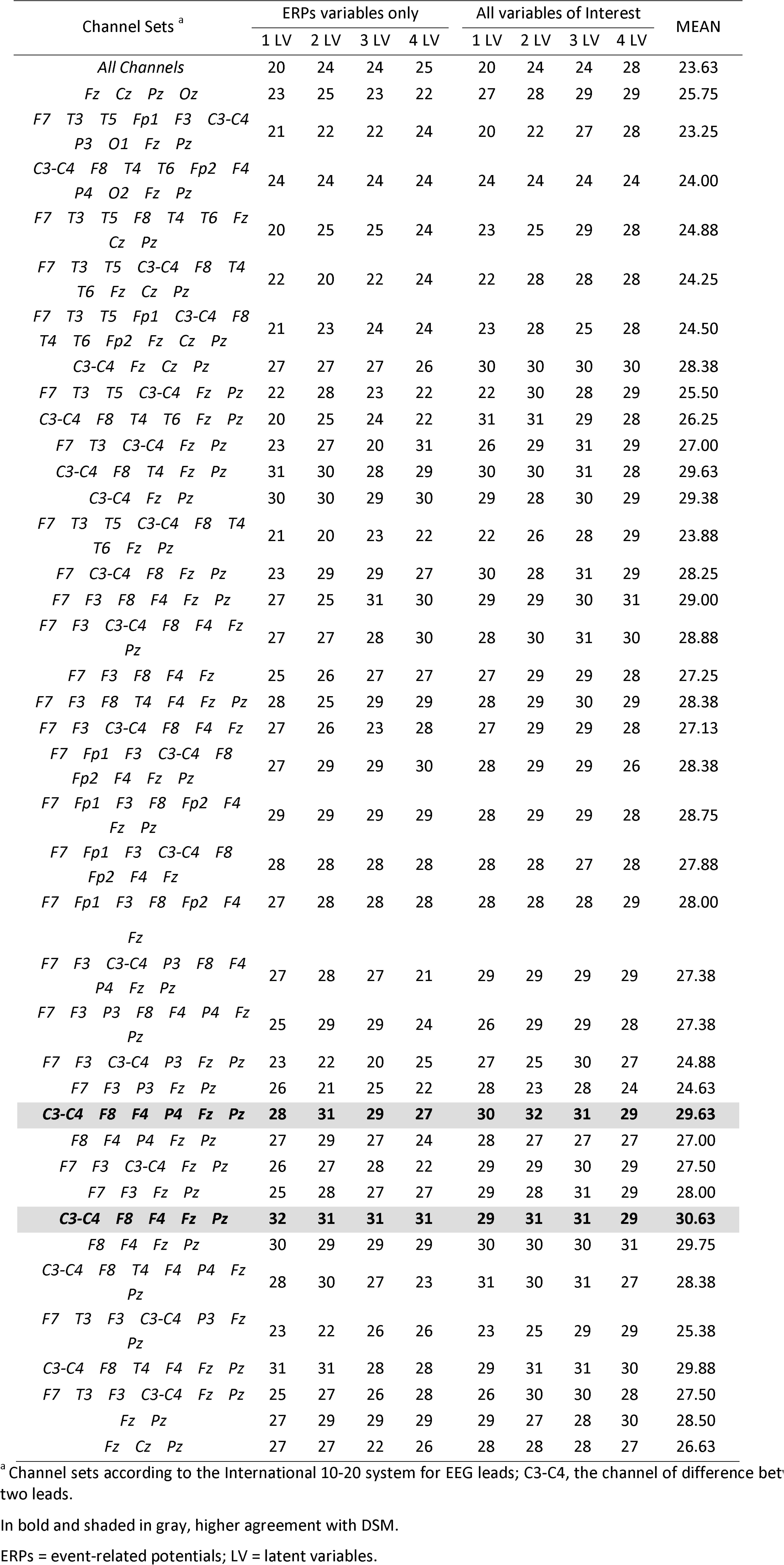
Agreement with DSM (Number of Subjects) by Channel Set and Number of Latent Vectors for Variables of Interest

The data from each investigated set of EEG channels were analyzed either with or without behavioral (ANT) and psychological (WISC) variables. They were submitted to an unsupervised method called Clustering of Variables around Latent Variables (CLV) [17,18]. The aim was to reduce the number of dimensions in the variables space by extracting latent variables (LVs), each of them associated with a cluster of the VOI variables. This clustering approach is based on correlation similarity indices between variables and aims to identify directional clusters of variables. As it is, each extracted LV, associated with a specific cluster of variables, is defined as the first (standardized) principal component of the variables assigned to that cluster [21]. In our study, the clusters of variables around one to four LVs were systematically considered and the best number of LVs was determined according to the classification of subjects.

In fact, our ultimate purpose was to reindex the subjects into two groups according to the objective behavioral, psychological, and neurophysiological data, all of which were summarized around the extracted LVs. A k-means clustering method [27] was performed on the subjects using the LVs. Each reclassification obtained (named “output”) was checked for agreement compared to the classification of subjects using the DSM-IV-TR (called “DSM agreement”). The probability that agreement with DSM could be observed under randomness was checked using the Chi-Square test for categorical variables with one degree of freedom, and the expected value was 20 (absolute randomness). The kappa index [28] was also evaluated to estimate the agreement level with DSM (S2 method).

In order to determine whether reclassification was consistent, this study analyzed which subjects were reclassified into another group other than his/her original DSM group (“miss-reclassification”). This was performed for each specified agreement rate with DSM, based on the outputs that provided this level of agreement. A measure of “classifier power” was suggested by dividing the level of agreement with DSM and the number of miss-reclassified subjects in any output with that agreement with DSM.

Subsequently, the outputs with the highest classifier power were considered the most representative to estimate the validity of the DSM. Among all outputs, these were submitted to cross-validation using the Leave-One-Out (LOO) strategy, where the consistency of agreement with DSM *versus* LV was observed across the mean values of the 39 LOO outputs, with 38 subjects each one. The output with the highest consistency of agreement with DSM was regarded as the most representative.

Using the most representative outputs, a sensitivity and specificity analysis was suggested for DSM criteria for ADHD.

The dataset with variables used in the multivariate analysis is available in the supplementary material (S3 Dataset).

## Results

### Evaluating predictor and confounding variables

Control and ADHD groups showed significant differences in the quantitative scores of DSM-IV-TR (p < 0.05) (Table 1). The significance level of the difference between the two groups was higher for the inattention score than for the hyperactivity/impulsivity score. Among variables considered as confounders, the I.Q. scores in ADHD group were lower than in TD boys (p = 0.006) although they were always higher than 80. All other variables, including learning rate, were not significantly different between the two groups according to the Student’s t-test (normality and homoscedasticity were not statistically rejected).

### Overview of Event-Related Potentials

Cue and target-related potentials with late peak latency (> 200ms, corresponding to parietal P3) were observed in all channels. Peak amplitude varied from the maximum value of 18μV for the frontopolar target ERP to the minimum value of 1μV for the difference between C3 and C4 (Fig 1).

### Data Mining and Reclassification of subjects

Values of agreement with DSM ranging from 0.51 (20 subjects, p = 0.125, χ^2^ = 0.00, p = 1.000) to 0.82 (32 subjects, χ^2^ = 7.20, p = 0.004) were found for all 40 channel sets and LV sets (extracting 1 to 4 LVs), with or without behavioral variables enclosed (Table 2). The *worst* mean agreement level between DSM and K-means reclassification was 23.25 for [F7, T3, T5, Fp1, F3, C3-C4, P3, O1, Fz, Pz] channel set (χ^2^ = 0.43, p = 0.528); and the best mean agreement level with DSM was 30.62 for [C3-C4, F8, F4, Fz, Pz] channel set (χ^2^ = 5.63, p = 0.010).

The two channel sets that manifested the highest agreement with DSM were [C3-C4, F8, F4, Fz, Pz] (using 1 LV from ERP variables alone) and [C3-C4, F8, F4, P4, Fz, Pz] (using 3 LVs from ERP and behavioral variables together). Table 2 shows all the observed agreement rates for all channel sets.

As expected, there was an inverse relationship between the agreements with DSM (from 20 to 32, according to the trial) and the total number of miss-reclassified subjects (Fig 2). The highest agreement rate found (32 subjects) corresponded to the highest classifier power (4.00).

**Figure 2.**
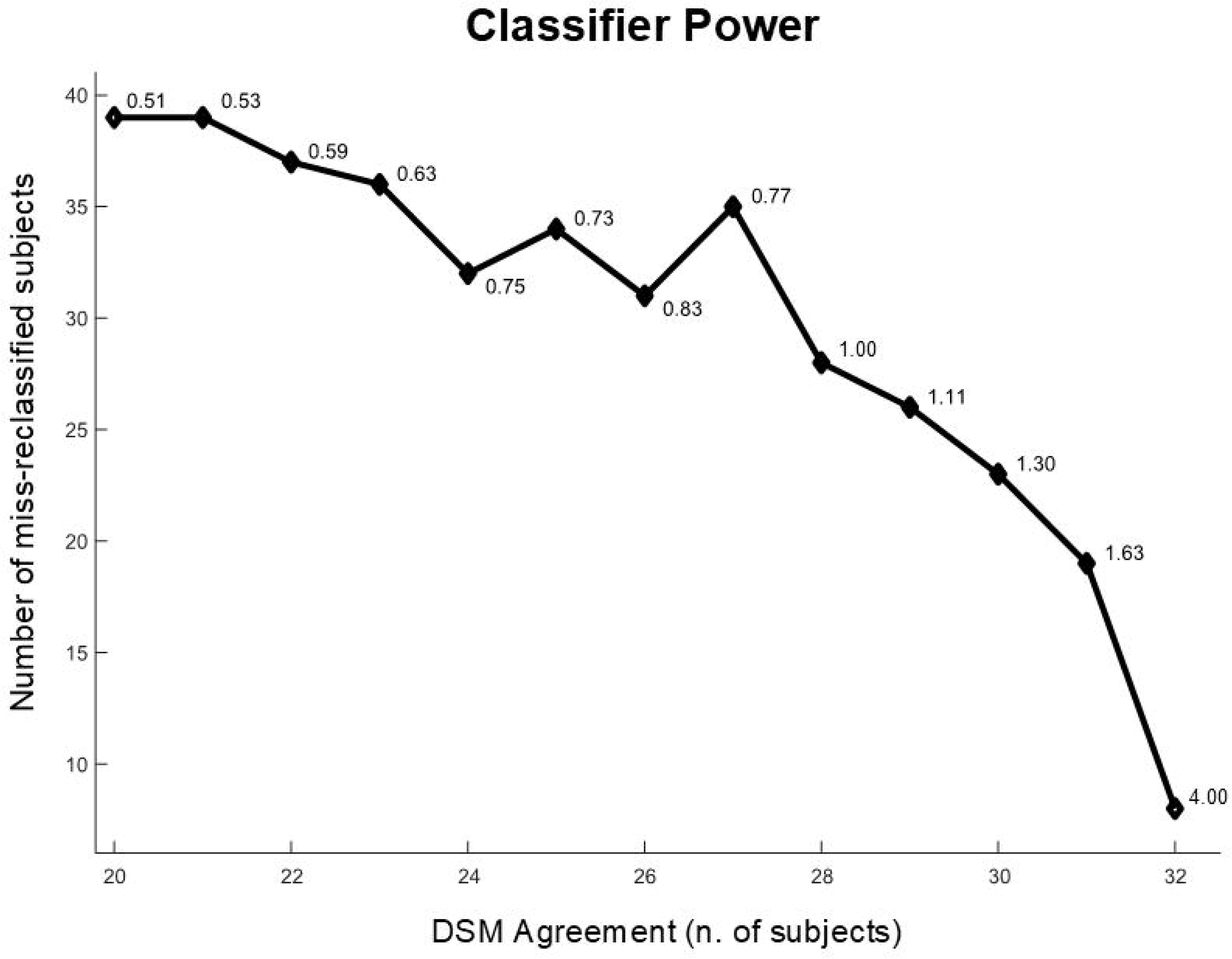
Classifier Power of the MVA analysis. Number of miss-reclassified subjects per agreement with DSM (number of subjects) in all 320 outputs, and the corresponding classifier power (numbers at the graph marks). See methods.

Regarding an agreement level equal to 32, the LOO cross-validation showed that only the output from the [C3-C4, F8, F4, Fz, Pz] channel set (using 1 LV from ERP variables alone) was valid, and therefore, it was the most representative: the mean agreement of all 39 outputs was 31.56 subjects (the highest mean agreement among all others, see S4 Output), while the output from [C3-C4, F8, F4, P4, Fz, Pz] (using 3 LVs from ERP and behavioral variables together) had a mean agreement equal to 29.87 subjects. The S4 Output has the agreement data of all outputs and their respective LOO cross-validations (mean values of all 39 outputs).

Seven subjects in the selected output ([C3-C4, F8, F4, Fz, Pz] channel set, using 1 LV from ERP variables alone) were miss-reclassified: C005, C009, C012, C020, C023 (control subjects miss-reclassified to the ADHD group) and T010, T026 (ADHD subjects miss-reclassified to the Control group). Based on these results, the estimated specificity and sensitivity of the DSM-IVTR regarding biological variables were 75.00% and 89.47%, respectively, with accuracy of 82.05% corresponding to the kappa index of 0.75, i.e. *a substantial strength of agreement* [28].

## Discussion

Our study tackled the possibility of examining attention deficit/hyperactivity disorder using the DSM manual, whose criteria seemed to be biologically justified for ADHD diagnosis with “substantial strength of agreement” regarding a specific pool of behavioral, psychological, and neurophysiological variables. A larger source of behavioral and neurophysiological data related to ADHD phenomenology was found in the ANT, since this test is suitable to access the three dimensions of attention in the same setting [22, 29].

In order to classify subjects as normal or ADHD according to DSM criteria, caregivers were asked which criteria were well-defined characteristics of their children. Other assessment methodologies should also be used along with DSM for ADHD diagnosis both in scientific research and in clinical practice. Moreover, the manual should not be applied as a questionnaire in clinical practice. However, in this case, the object of evaluation is the DSM itself; it was thus the only clinical diagnostic tool used for the classification of subjects as either ADHD or not-ADHD. It was the most objective and controlled way to obtain the scores without interference from any other clinical impression. Supporting the present methodology, the ASRS (the WHO questionnaire for adult ADHD screening) was self-administered [30].

The most informative neurophysiological variables in this pool proved to be topographically asymmetrical and represented the binded information from ERPs in the three time intervals of interest from mid-frontal, mid-parietal, right frontal, and right anterior temporal sites (Fz, Pz, F4, and F8 channels), as well as early ERP components (45 to 290ms after target onset) from the ‘C3 minus C4’ channel. This asymmetry has already been shown to correlate with DSM scores, mainly with hyperactivity/impulsivity [31]. It was not the focus of the present investigation to establish models for ADHD or its neural mechanisms. However, our results revealed certain biological aspects of ADHD regarding alterations in the frontal control over executive functions [21, 32, 33]. Several studies have shown brain asymmetries related to ADHD phenomenology, especially in the frontal-striatal network [33-35]. Asymmetrical topographic patterns were shown in our previous studies by spectral and coherence analyses of the resting EEG, with signs of relative inactivation of the frontal and *left* temporal cortex, known to be responsible for voluntary attention, and which are impaired in ADHD patients [36]. These observations could partially explain the leading role of ERP data from the right frontal regions in discriminating between patients and controls in the present study.

Our results suggest that DSM-IV-TR is actually an effective and biologically justified tool for ADHD diagnosis. Although a Kappa index of 0.75 should be considered with reservations for some clinical settings, a clinical method such as DSM, with sensitivity of 89.47% and specificity of 75% with regard to objective behavioral, psychological, and neurophysiological measures, should be considered as particularly relevant considering the complexity of mental disorders, especially those with dimensional clinical phenomenology and without any validated biomarkers, so much so that the only psychiatric tool for diagnosis and intervention is still phenomenological examination, which must be systematically (although qualitatively) performed based on manuals such as the DSM. Furthermore, the ADHD has been regarded as a heterogeneous condition with inattentive, hyperactive, and combined subtypes, which could be the answer for this level of agreement since these subtypes were not predicted for reclassification. The existence of three discrete subtypes could lead to a flawed reclassification.

The study was designed with a relatively small sample size. Any subject with the slightest suspicion of comorbidity was excluded. Potentially confounding factors commonly ignored were monitored (such as manipulating technologies that could lead to the development of specific skills that would interfere with test performance and socioeconomic niche that could determine psychological development). Therefore, the sample should be regarded as well consistent with and representative of ADHD. On the other hand, in order to classify the subjects according to their mental condition (a multidimensional phenomenological complex) it would seem reasonable to gather a considerable number of objective variables (as many as possible). Brain processing during the ANT test as well as ADHD neurophysiology and its behavioral correlates in this study were considered as potentially relevant features requiring further investigation. Since it did not seem feasible to collect a sample with thousands of subjects evaluating hundreds of variables in an experiment with optimal control of confounders and bias, the dataset had to have a much smaller number of observations than variables. A small but strictly controlled and well-exploited sample was considered preferable, even in conditions of high prevalence such as the ADHD. Multivariate statistical modeling, such as Linear Regression, Discriminant Analysis, or Factor Analysis, are well known as unsuitable for this situation. Thus, in order to tackle a frequently encountered methodological problem, often referred to as the curse of the dimensionality, an explorative data analysis suitable for small samples was adopted to identify a small number of synthetic latent variables. Instead of the Principal Component Analysis, which does not always provide easily interpretable latent variables (the principal components), an approach with clustering of variables was chosen to reduce data dimensionality to a small number of new latent variables. The CLV method [17,18] has already been applied to a wide range of research domains among which are pattern analysis [37], chemometrics [38], image analysis [39], and analysis scorecards in the health sector [40]. The results seemed promising for optimizing the experimental designs of clinical studies in which it is difficult to recruit large groups of patients.

The estimated I.Q. scores were different between the groups diagnosed using DSM-IV-TR, but this was not regarded as bias. Literature has shown that intelligence tests are sensitive to ADHD, and patient scores are generally lower than controls [41]. Even taking into account the above restrictions, there is no doubt that the data obtained seem to be quite consistent.

A major limitation of this study is the impossibility of testing all possible combinations among variables. Since the consistency of our paradigm lies on the fact that no biological / behavioral assumptions were considered in the grouping of variables, it is necessary to test them randomly grouped. However, there are more than one million possible combinations of EEG channels. The number of combinations grows exponentially if the other variables are included individually. For computational feasibility, sets of variables were arbitrarily chosen for analysis. Considering the eighty sets studied, it can be assumed that DSM has at least a “substantial level of agreement” with biological determinants. Perhaps some untested set of variables may manifest higher agreement with the DSM, but the results found already satisfy our objective of testing the validity of the DSM criteria for ADHD.

The aim of the present study was to draw attention to psychiatric approaches based on diverse and multiple factors intervening in the biological mechanisms of human behavior and its disorders. Therefore, we have found that there is a consistent complex of biological information, which is in accordance with previous knowledge on ADHD, that confers biological validity for DSM criteria for ADHD.

## Aknowledgements

We would like to thank The Child’s Neurology program of the National Institute of Women, Children, and Adolescents’ Health Fernandes Figueira. We are also grateful to Mrs. Maria S. Santana and Mrs. Aldenys Perez for the technical support. Our team is especially grateful to our friend, colleague, and co-author Dr. Adailton Tadeu Pontes, children’s neurologist, who greatly inspired this article, and passed away during its preparation.

---

**S1 table. Time intervals of Interest.** Windows holding Cue-related potentials (CUE), contingent voltage variation (CVV), and Target-related potentials (TGT) for each channel (ms), to calculate peak amplitude, peak latency, and mean amplitude. Zero ms refers to Target onset (see Figure 1).

**S2 method. Calculating the Kappa Index.** Inference of the magnitude of agreement between two categorical data (in this case, subjects’ classification according to DSM and to biological/behavioral variables in the dataset).

**S3 dataset. DSM scores and variables of interest.** *In the first sheet:* innatention, hyperactive+impulsive, and total scores for ADHD according to DSM-IV-TR; *second sheet:* all data concerning (1) scores of the WISC-III subtests, (2) ANT performance (AC: accuracy, RT: reaction time and IVRT: intraindividual variation in reaction time, which is the standard deviation of the subject’s RT) for each experimental condition (Allcon: mean value of all conditions, NtCue: neutral cue condition, SpCue: spatial cue condition, ConTg: congruent target condition, IncTg: incongruent target condition) and attentional network (alerting, orienting, and executive) plus learning indexes (IVRT or RT at the last ANT block/first ANT block), and (3) the neurophysiological measures (peak amplitudes and latencies, and mean amplitudes inside each time interval of interest in S1) for the EEG channel set.

**S4 output. DSM versus Biological agreement.** The channel sets (rows) and number of latent vectors extracted (rows) holding ERPs variables alone or ERPs + behavioral (ANT and WISC) variables. The first block (on the left) are outputs from all 39 subjects. The second block (on the right) are the averages of outputs with n-1 subjects (LOO cross-validation).

## References

1. American Psychiatric Association. Diagnostic and statistical manual of mental disorders. 4th ed. Washington, DC: Author; 1994.

2. American Psychiatric Association. Diagnostic and statistical manual of mental disorders. 5th ed. Arlington, VA: American Psychiatric Publishing; 2013.

3. Insel T. Director’s Blog: Transforming Diagnosis. 2013 Apr 29 [cited 06 Jul 2018] In: The National Institute of Mental Health Website. [about 2 screens] Available from: http://www.nimh.nih.gov/about/director/2013/transforming-diagnosis.shtml.

4. Wakefield JC. DSM-5, psychiatric epidemiology and the false positives problem. Epidemiol Psychiatr Sci. 2015 Jun;24(3):188–96. doi: 10.1017/S2045796015000116.

5. Rafalovich A. Exploring clinician uncertainty in the diagnosis and treatment of attention deficit hyperactivity disorder.Sociol Health Illn. 2005 Apr;27(3):305–23.

6. Strauss V. An ADHD controversy in the mental health community. 2013 February 13 [cited 06 July 2018]. In: The Washington Post [Internet]. Available from: https://www.washingtonpost.com/blogs/answer-sheet/post/an-adhd-controversy-in-the-mental-health-community/2012/02/12/gIQAHJun9Q_blog.html.

7. Maser JD, Akiskal HS. Spectrum concepts in major mental disorders. Psychiatr Clin North Am. 2002 Dec;25(4):xi–xiii.

8. Kirmayer LJ, Crafa D. What kind of science for psychiatry? Front Hum Neurosci. 2014 Jun 20;8:435. doi: 10.3389/fnhum.2014.00435.

9. Furman LM. Attention-deficit hyperactivity disorder (ADHD): does new research support old concepts? J Child Neurol. 2008 Jul;23(7):775–84. doi: 10.1177/0883073808318059.

10. Freitas-Silva LR, Ortega F. Biological determination of mental disorders: a discussion based on recent hypotheses from neuroscience. Cad Saude Publica. 2016 Aug 29;32(8):e00168115. doi: 10.1590/0102-311X00168115.

11. Scarr E, Millan MJ, Bahn S, Bertolino A, Turck CW, Kapur S, Möller HJ, Dean B. Biomarkers for Psychiatry: The Journey from Fantasy to Fact, a Report of the 2013 CINP Think Tank. Int J Neuropsychopharmacol. 2015 Apr 21;18(10):pyv042. doi: 10.1093/ijnp/pyv042.

12. Bochkarev VK, Lazarev VV, Nikiforov AI, Paniushkina SV, Severnyĭ AA. Clinico-electroencephalographic correlations in asthenoadynamic subdepressions.Zh Nevropatol Psikhiatr Im S S Korsakova. 1987;87(4):564–70.

13. Lazarev VV. The relationship of theory and methodology in EEG studies of mental activity. Int J Psychophysiol. 2006 Dec;62(3):384–93. Epub 2006 Mar 10.

14. Lazarev VV, Lebedeva IS, Tsutsul’kovskaia MIa, Severnyĭ AA, Pozharitskaia DA. Specificity of electroencephalographic correlations of asthenic disorders in atypical juvenile asthenic-depressive conditions. Zh Nevropatol Psikhiatr Im S S Korsakova. 1990;90(10):73–7.

15. Monakhov K, Perris C, Botskarev VK, von Knorring L, Nikiforov AI. Functional interhemispheric differences in relation to various psychopathological components of the depressive syndromes. A pilot international study.Neuropsychobiology. 1979;5(3):143–55.

16. Schwarz E, Izmailov R, Spain M, Barnes A, Mapes JP, Guest PC, Rahmoune H, Pietsch S, Leweke FM, Rothermundt M, Steiner J, Koethe D, Kranaster L, Ohrmann P, Suslow T, Levin Y, Bogerts B, van Beveren NJ, McAllister G, Weber N, Niebuhr D, Cowan D, Yolken RH, Bahn S. Validation of a blood-based laboratory test to aid in the confirmation of a diagnosis of schizophrenia. Biomark Insights. 2010 May 12;5:39–47.

17. Vigneau E, Qannari EM. Clustering of Variables Around Latent Components. Communications in Statistics - Simulation and Computation. 2003; 32: 1131–1150.

18. Vigneau E, Chen M, Qannari E M. ClustVarLV: An R Package for the Clustering of Variables Around Latent Variables. The R Journal. 2015; 7: 134–148.

19. Matuschek T, Jaeger S, Stadelmann S, Dölling K, Grunewald M, Weis S, von Klitzing K, Döhnert M. Implementing the K-SADS-PL as a standard diagnostic tool: Effects on clinical diagnoses. Psychiatry Res. 2016 Feb 28;236:119–24. doi: 10.1016/j.psychres.2015.12.021.

20. Hruby T, Marsalek P. Event-related potentials‐‐the P3 wave. Acta Neurobiol Exp (Wars). 2003;63(1):55–63.

21. Barry RJ, Johnstone SJ, Clarke AR. A review of electrophysiology in attentiondeficit/hyperactivity disorder: II. Event-related potentials. Clin Neurophysiol. 2003 Feb;114(2):184–98.

22. Fan J, McCandliss BD, Sommer T, Raz A, Posner MI. Testing the efficiency and independence of attentional networks. J Cogn Neurosci. 2002 Apr 1;14(3):340–7.

23. Kratz O, Studer P, Malcherek S, Erbe K, Moll GH, Heinrich H. Attentional processes in children with ADHD: an event-related potential study using the attention network test. Int J Psychophysiol. 2011 Aug;81(2):82–90. doi: 10.1016/j.ijpsycho.2011.05.008.

24. Wechsler D. Wechsler Intelligence Scale for Children(WISC-III): Manual. 3th ed. San Antonio: The Psychological Corporation; 1991.

25. Mello CB, Argollo Shayer PBM, Abreu N, AbreuII N, GodinhoIII K, Durán P, Vargem F, Muskat M, Miranda MC, Bueno OFA. Abbreviated version of the WISC-III: correlation between estimated IQ and global IQ of Brazilian children, Psicologia: Teor e Pesq. 2011; 27 (2): 149–155. dx.doi.org/10.1590/S0102-37722011000200002.

26. Abramov DM, Pontes M, Pontes AT, Mourao-Junior CA, Vieira J, Quero Cunha C, Tamborino T, Galhanone PR, deAzevedo LC, Lazarev VV. Visuospatial information processing load and the ratio between parietal cue and target P3 amplitudes in the Attentional Network Test. Neurosci Lett. 2017 Apr 24;647:91–96. doi: 10.1016/j.neulet.2017.03.031.

27. Gore Jr PA. Cluster Analysis, In: Tinsley HEA, Brown SD, editors. Handbook of Applied Multivariate Statistics and Mathematical Modeling. San Diego, CA: Academic Press; 2000. p. 297–321.

28. Landis JR, Koch GG. The measurement of observer agreement for categorical data. Biometrics. 1977 Mar;33(1):159–74.

29. Petersen SE, Posner MI. The attention system of the human brain: 20 years after. Annu Rev Neurosci. 2012;35:73–89. doi: 10.1146/annurev-neuro-062111-150525.

30. Kessler RC, Adler L, Ames M, Demler O, Faraone S, Hiripi E, Howes MJ, Jin R, Secnik K, Spencer T, Ustun TB, Walters EE. The World Health Organization Adult ADHD Self-Report Scale (ASRS): a short screening scale for use in the general population. Psychol Med. 2005 Feb;35(2):245–56.

31. Abramov DM., Cunha CQ, Pontes MC, Galhanone PR, deAzevedo LC, Lazarev VV. Functional Asymmetry In The Central Brain Regions In Boys With Attention Deficit Hyperactivity Disorder Detected By Event Related Potentials During Performance Of The Attentional Network Test. 2016. Preprint.Avaiable on BioRxiv, https://doi.org/10.1101/118380; cited on 06 July 2018.

32. Barkley RA. Behavioral inhibition, sustained attention, and executive functions: constructing a unifying theory of ADHD.Psychol Bull. 1997 Jan;121(1):65–94. doi: 10.1037/0033-2909.121.1.65.

33. Barkley RA. Attention-Deficit/Hyperactivity Disorder: handbook for diagnosis and treatment. 3th ed. New York: The Guilford Press; 2006. p 32–33.

34. Langleben DD, Austin G, Krikorian G, Ridlehuber HW, Goris ML, Strauss HW. Interhemispheric asymmetry of regional cerebral blood flow in prepubescent boys with attention deficit hyperactivity disorder. Nucl Med Commun. 2001 Dec;22(12): 1333–40.

35. Dang LC, Samanez-Larkin GR, Young JS, Cowan RL, Kessler RM, Zald DH. Caudate asymmetry is related to attentional impulsivity and an objective measure of ADHD-like attentional problems in healthy adults. Brain Struct Funct. 2016 Jan;221(1):277–86. doi: 10.1007/s00429-014-0906-6.

36. Lazarev VV, Pontes M, Pontes AT, Vieira J, Quero-Cunha C, Tamborino T, deAzevedo LC, Abramov DM. EEG and ERP characteristics of Attention Deficit Hyperactivity Disorder in children and adolescents [abstract]. Int. J. Psychophysiol. 2106; 108: 75.

37. Vigneau E, Sahmer K, Qannari EM, Bertrand D. Clustering of variables to analyze spectral data. J Chemometrics, 2005; 19: 122–128.

38. Vigneau E, Qannari EM, Navez B, Cottet V. Segmentation of consumers in preference studies while setting aside atypical or irrelevant consumers. Food Quality Pref. 2005; 47: 54–63.

39. Legland D, Beaugrand J. Automated clustering of lignocellulosic fibres based on morphometric features and using clustering of variables. Industrial Crops and Products; 2013; 45: 253–261.

40. Lovaglio PG. Model building and estimation strategies for implementing the Balanced Scorecard in Health sector. Quality & Quantity. 2011; 45: 199–212.

41. Mackenzie GB, Wonders E. Rethinking Intelligence Quotient Exclusion Criteria Practices in the Study of Attention Deficit Hyperactivity Disorder. Front Psychol. 2016 May 31;7:794. doi: 10.3389/fpsyg.2016.00794.

